# Information-dense transcription factor binding site clusters identify target genes with similar tissue-wide expression profiles and serve as a buffer against mutations

**DOI:** 10.1101/283267

**Authors:** Ruipeng Lu, Peter K. Rogan

## Abstract

**Background:** The distribution and composition of *cis*-regulatory modules (e.g. transcription factor binding site(TFBS) clusters) in promoters substantially determine gene expression patterns and TF targets, whose expression levels are significantly regulated by TF binding. TF knockdown experiments have revealed correlations between TF binding profiles and gene expression levels. We present a general framework capable of predicting genes with similar tissue-wide expression patterns from activated or repressed TF targets using machine learning to combine TF binding and epigenetic features.

**Methods:** Genes with correlated expression patterns across 53 tissues were identified according to their Bray-Curtis similarity. DNase I-accessible promoter intervals of direct TF target genes were scanned with previously derived information theory-based position weight matrices (iPWMs) of 82 TFs. Features from information density-based TFBS clusters were used to predict target genes with machine learning classifiers. The accuracy, specificity and sensitivity of the classifiers were determined for different feature sets. Mutations in TFBSs were also introduced to *in silico* examine their impact on cluster densities and the regulatory states of target genes.

**Results:** We initially chose the glucocorticoid receptor gene (*NR3C1*), whose regulation has been extensively studied, to test this approach. *SLC25A32* and *TANK* were found to exhibit the most similar expression patterns to this gene across 53 tissues. A Decision Tree classifier exhibited the largest area (0.9987 and 0.9956 respectively before and after eliminating inaccessible promoter intervals based on DNase I HyperSensitive Regions (DHSs)) under the Receiver Operating Characteristic (ROC) curve in detecting such coordinately regulated genes. Target gene prediction was confirmed using siRNA knockdown data of TFs, which was more accurate than CRISPR inactivation. *In-silico* mutation analyses of TFBSs also revealed that one or more information-dense TFBS clusters in promoters are required for accurate target gene prediction.

**Conclusions:** Machine learning based on TFBS information density, organization, and chromatin accessibility accurately identifies gene targets with comparable tissue-wide expression patterns. Multiple, information-dense TFBS clusters in promoters appear to protect promoters from the effects of deleterious binding site mutations in a single TFBS that would effectively alter the expression state of these genes.

## INTRODUCTION

The distinctive organization and combination of transcription factor binding sites (TFBSs) and regulatory modules in promoters dictates specific expression patterns within a set of genes [1]. Clustering of multiple adjacent binding sites for the same TF (homotypic clusters) and for different TFs (heterotypic clusters) defines *cis*-regulatory modules (CRMs) in human gene promoters and can amplify the influence of individual TFBSs on gene expression through increasing binding affinities, facilitated diffusion mechanisms and funnel effects [2]. Because tissue-specific TF-TF interactions in TFBS clusters are prevalent, these features can assist in identifying correct target genes by altering binding specificities of individual TFs [3]. Previously, we derived iPWMs from ChIP-seq data that can accurately detect TFBSs and quantify their strengths by computing associated *R*_*i*_ values (Rate of Shannon information transmission for an individual sequence [4]), with *R*_*sequence*_ being the average of *R*_*i*_ values of all binding site sequences and representing the average binding strength of the TF [3]. Furthermore, information density-based clustering (IDBC) can effectively identify functional TF clusters by taking into account both the spatial organization (i.e. intersite distances) and information density (i.e. *R*_*i*_ values) of TFBSs [5].

TF binding profiles, either derived from *in vivo* ChIP-seq peaks [6–8] or computationally detected binding sites and CRMs [9], have been shown to be predictive of absolute gene expression levels using a variety of tissue-specific machine learning classifiers and regression models. Because signal strengths of ChIP-seq peaks are not strictly proportional to TFBS strengths [3], representing TF binding strengths by ChIP-seq signals may not be appropriate; nevertheless, both achieved similar accuracy [10]. CRMs have been formed by combining two or three adjacent TFBSs [9], which is inflexible, as it arbitrarily limits the number of binding sites contained in a module, and does not consider differences between information densities of different CRMs. Chromatin structure (e.g. histone modification (HM) and DNase I hypersensitivity) were also found to be statistically redundant with TF binding in explaining tissue-specific mRNA transcript abundance at a genome-wide level [7,8,11,12], which was attributed to the heterogeneous distribution of HMs across chromatin domains [8]. Combining these two types of data explained the largest fraction of variance in gene expression levels in multiple cell lines [7,8], suggesting that either contributes unique information to gene expression that cannot be compensated for by the other.

The number of genes directly bound by a TF significantly exceeds the number of differentially expressed (DE) genes whose expression levels significantly change upon knockdown of the TF. Only a small subset of direct target genes whose promoters overlap ChIP-seq peaks were DE after individually knocking 59 TFs down using small interfering RNAs (siRNAs) in the GM19238 cell line [13]. Using these knockdown data on 8,872 genes as the gold standard, correlation between TFBS counts and gene expression levels across 10 different cell lines was more predictive of DE targets than setting a minimum threshold on TFBS counts [14]. Their TFBS counts were defined as the number of ChIP-seq peaks overlapping the promoter, though it was unknown how many binding sites were present in these peaks; positives might not be direct targets in the TF regulatory cascade, as the promoters of these targets were not intersected with ChIP-seq peaks. By perturbing gene expression with CAS9-directed clustered regularly interspaced short palindromic repeats (CRISPR) of 10 different TF genes in K562 cells, the regulatory effects of each TF on 22,046 genes were dissected by single cell RNA sequencing with a regularized linear computational model [15]; this accurately revealed DE targets and new functions of individual TFs, some of which were likely regulated through direct interactions at TFBSs in their corresponding promoters. Machine learning classifiers have also been applied in a small number of gene instances to predict targets of a single TF using features extracted from *n*-grams derived from consensus binding sequences [16], or from TFBSs and homotypic binding site clusters [5].

To investigate whether the distribution and composition of CRMs in promoters substantially determines gene expression profiles of direct TF targets, we developed a general machine learning framework that predicts which genes have similar expression profiles to a given gene and predicts DE direct TF targets by combining information theory-based TF binding profiles with DHSs. Upon filtering for accessible promoter intervals with DHSs, features designed to capture the spatial distribution and information composition of CRMs were extracted from clusters identified by the IDBC algorithm from iPWM-detected TFBSs. Though not all direct targets regulated by multiple TFs share a common tissue-wide expression profile, this framework provides insight into the transcriptional program of genes with similar profiles by dissecting their *cis*-regulatory element organization and strengths. We identify genes with comparable tissue-wide expression profiles by application of Bray-Curtis similarity [17]. Using transcriptome data generated by CRISPR-[15] and siRNA-based [13] TF knockdowns, we predicted DE TF target genes that are simultaneously direct targets whose promoters overlap tissue-specific ChIP-seq peaks.

## MATERIALS AND METHODS

To identify genes with similar tissue-wide expression patterns, we formally define gene expression profiles and pairwise similarity measures between profiles of different genes. A general machine learning framework relates features extracted from the organization of TFBSs in these genes to their tissue-wide expression patterns. Since protein-coding (PC) sequences represent the most widely studied and best understood component of the human genome [18], positives and negatives for predicting DE direct TF target genes that encode proteins (TF targets for short below) were obtained from CRISPR- and siRNA-generated knockdown data (see below).

### Similarity between gene expression profiles

For each of 56,238 genes, the Genotype-Tissue Expression (GTEx) project measured its expression levels in 53 tissues in a number of individuals (*N*=5-564), and provides the median expression value (in RPKM (Reads Per Kilobase of transcript per Million mapped reads) in the GTEx Analysis v6p release) of each gene in each tissue [19]. To capture the tissue-wide overall expression pattern of a gene instead of within a single tissue, the expression profile of a gene was defined as its median RPKM across the 53 tissues. In Equation 1, the expression profile of a gene is described by a vector of size 53; note that different isoforms whose expression patterns may significantly differ from each other cannot distinguished by this approach.

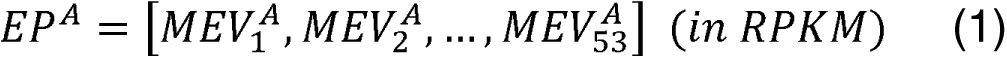

Where *EP*^*A*^ is the expression profile of Gene *A*, 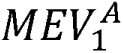 is the median expression value of Gene *A* in Tissue 1, 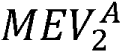 is the median expression value of Gene *A*, in Tissue 2, etc.

To discover other genes whose expression profiles are similar to a given gene, we computed the Bray-Curtis Similarity (Equation 2) between the expression profiles of gene pairs. Compared to other similarity metrics (Table 1, Example 1, Additional file 1), the application of this function is justified by desirable properties, including: 1) maintaining bounds between 0 and 1, 2) achieving the maximal similarity value 1 if and only if two vectors are identical, and 3) larger values having a larger impact on the resultant similarity value.

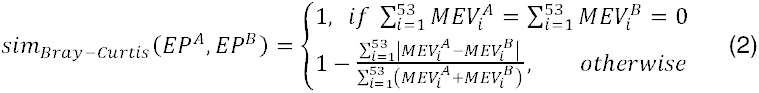

**Table 1:**
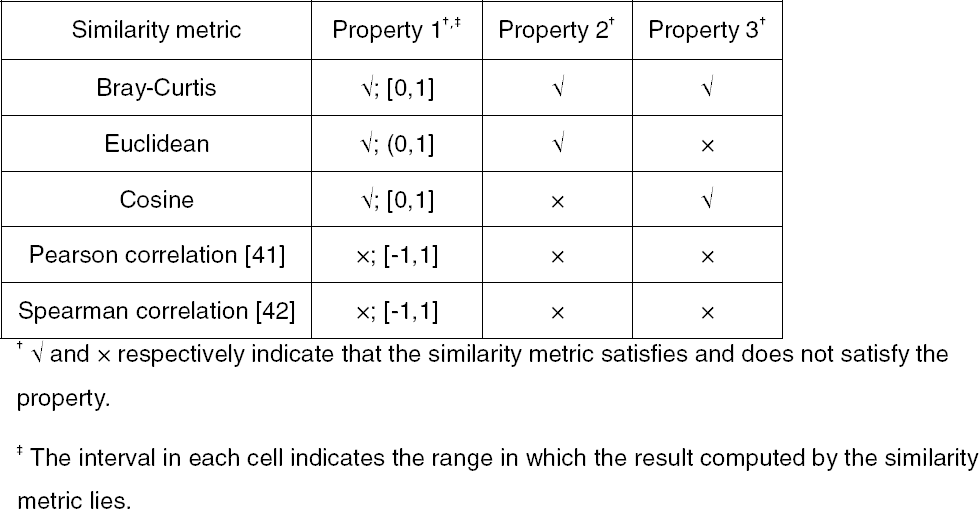
Comparison between metrics in measurement of similarity between gene expression profiles.

**Example 1**. Assume that Genes *A,B,C,D,E,F* respectively have the following expression profiles across two tissues: *EP*^*A*^ = [1,1], *EP*^*B*^ = [2,2], *EP*^*C*^ = [3,3], *EP*^*D*^ = [1,2], *EP*^*E*^ = [1,99], *EP*^*F*^ = [1,100]. The ground-truth similarity relationships that we can intuitively infer include *sim(EP*^*C*^*, EP*^*A*^)< *sim(EP*^*C*^*, EP*^*B*^)< 1, and *(EP*^*A*^*, EP*^*D*^)< *sim (EP*^*E*^*, EP*^*F*^)< 1. Only the results computed by the Bray-Curtis Similarity are completely concordant with these ground-truth relationships (Table 2).

**Table 2:** Similarity values computed by different metrics.

| Similarity metric | $sim(EP^C, EP^B)$ | $sim(EP^C, EP^A)$ | $sim(EP^E, EP^F)$ | $sim(EP^A, EP^D)$ |
| --- | --- | --- | --- | --- |
| Bray-Curtis | 0.8 | 0.5 | $\approx 0.995$ | 0.8 |
| Euclidean | $\approx 0.41$ | $\approx 0.26$ | 0.5 | 0.5 |
| Cosine | 1 | 1 | $\approx 0.999999995$ | $\approx 0.949$ |
| Pearson correlation | Undefined | Undefined | 1 | 1 |
| Spearman correlation | 1 | 1 | 1 | 1 |

### Prediction of genes with similar expression profiles

The framework for identifying genes whose expression profiles most resemble a particular gene is shown in Figure 1A and B. The genomic locations of all DHSs in 95 cell types generated by the ENCODE project [18; hg38 assembly] were selected for known promoters [21], then 94 iPWMs exhibiting primary binding motifs for 82 TFs [3] were used to detect TFBSs within the overlapping intervals. When detecting heterotypic TFBS clusters with the IDBC algorithm, a minimum threshold 0.1 \**R*_*sequence*_ was specified for the *R*_*i*_ values of TFBSs in order to remove overlapping intervals. When detecting heterotypic TFBS clusters with the IDBC algorithm, a weak binding sites that were more likely to be false positive TFBSs [3].

**Figure 1:**
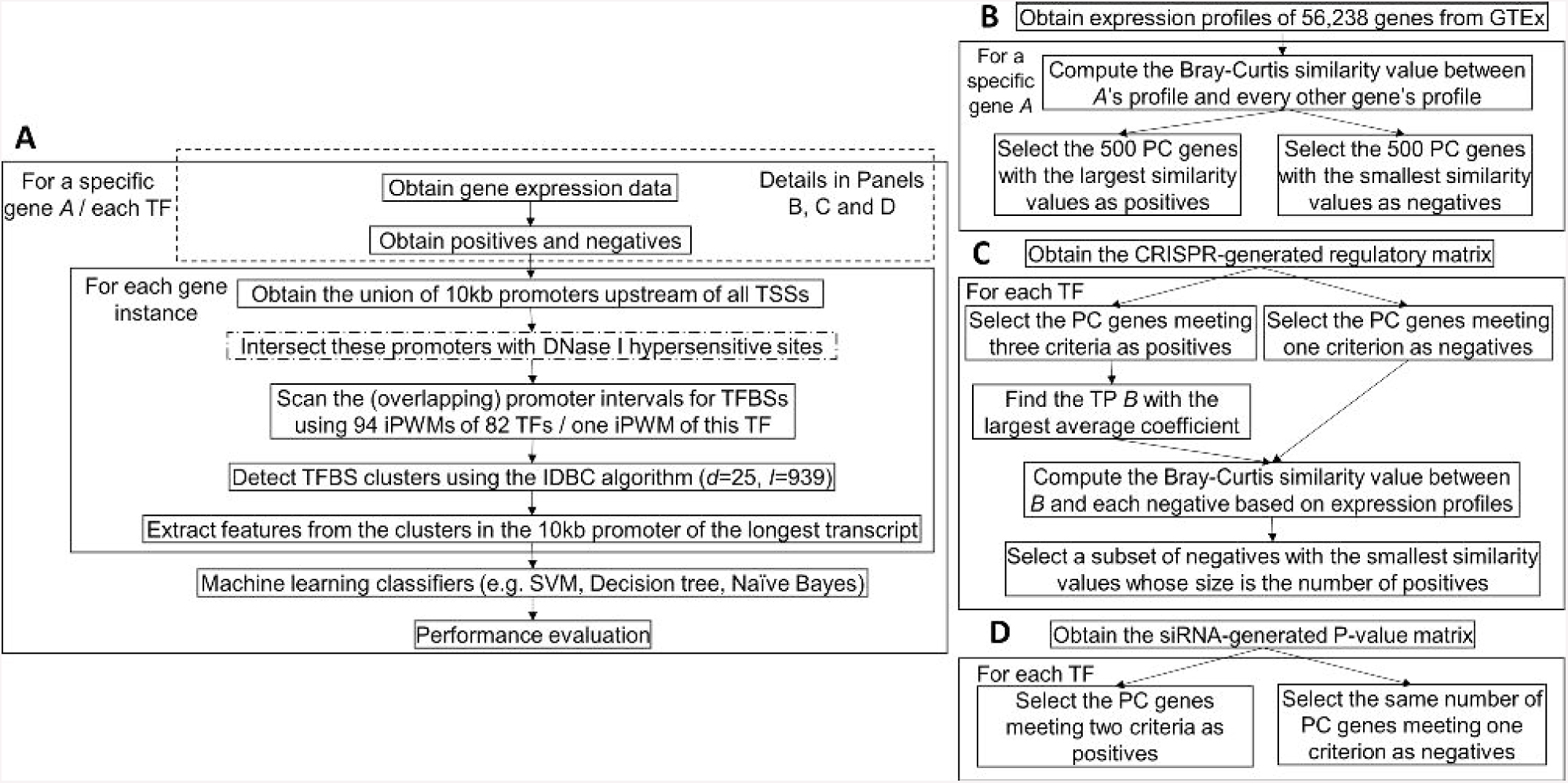
The general framework for predicting genes with similar tissue-wide expression profiles and TF targets. An overview of the machine learning framework. The steps enclosed in the dashed rectangle vary across prediction of genes with similar expression profiles and TF targets. The step with a dash-dotted border that intersects promoters with DHSs is a variant of the primary approach. In the IDBC algorithm (Additional file 1), the parameter *I* is the minimum threshold on the total information contents of TFBS clusters. In prediction of genes with similar expression profiles, the minimum value was 939, which was the sum of mean information contents (*R*_*sequence*_ values) of all 94 iPWMs; in prediction of direct targets, this value was the *R*_*sequence*_ value of the single iPWM used to detect TFBSs in each promoter. The parameter *d* is the radius of initial clusters in base pairs, whose value, 25, was determined empirically. The performance of seven different classifiers were evaluated with statistics (accuracy, sensitivity and specificity) (Additional file 1). **B)** Obtaining of the positives and negatives for identifying genes with similar expression profiles to a given gene (Additional file 2). **C)** Obtaining of the positives and negatives for predicting target genes of seven TFs using the CRISPR-generated perturbation data in K562 cells (Additional file 3). **D)** Obtaining of the positives and negatives for predicting target genes of 11 TFs using the siRNA-generated knockdown data in GM19238 cells (Additional file 4).

The information density-related features derived from each TFBS cluster included: 1) The distance between this cluster and the transcription start site (TSS); 2) The length of this cluster; 3) The information content of this cluster (i.e. the sum of *R*_*i*_ values of all TFBSs in this cluster); 4) The number of binding sites of each TF within this cluster; 5) The number of strong binding sites (*R*_*i*_ > *R*_*sequence*_) of each TF within this cluster; 6) The sum of *R*_*i*_ values of binding sites of each TF within this cluster; 7) The sum of *R*_*i*_ values of strong binding sites (*R*_*i*_ > *R*_*sequence*_) of each TF within this cluster.

For a gene, each of Features 1-3 was defined as a vector whose size equals the number of clusters in the promoter; thus, the entire vector could be input into a classifier. If two genes contained different numbers of clusters, the maximum number of clusters among all instances was determined, and null clusters were added at the 5’ end of promoters with fewer clusters, enabling all instances to have the same cluster count. Machine learning classifiers with default parameter values in MATLAB were used to generate ROC curves (Additional file 1).

### Prediction of TF targets

#### Using gene expression in the CRISPR-based perturbations

Dixit et al. performed CRISPR-based perturbation experiments using multiple guide RNAs for each of ten TFs in K562 cells, resulting in a regulatory matrix of coefficients that indicate the effect of each guide RNA on each of 22,046 genes [15]. The coefficient of a guide RNA on a gene is defined as the log_10_ (fold change in gene expression level) [15]. Among these ten TFs, we have previously derived iPWMs exhibiting primary binding motifs for seven (EGR1, ELF1, ELK1, ETS1, GABPA, IRF1, YY1) [3]. Therefore, the framework for predicting TF targets in the K562 cell line (Figure 1A and 1C) was applied to these TFs. The criteria for defining a positive (i.e. a target gene), of a TF were:

1. The fold change in the expression level of this gene for each guide RNA of the TF was > (or <) 1, consistent with the possibility that the gene was regulated by the TF, and
2. The average fold change in the expression level of this gene for all guide RNAs of the TF was > threshold, (or <1 / *ε*), and
3. The promoter interval (10 kb) upstream of a TSS of this gene overlaps a ChIP-seq peak of the TF in the K562 cell line.

If the coefficients of all guide RNAs of the TF for a gene are zero, the gene was defined as a negative. As the threshold *ε* increases, the number of positives strictly decreases; as *ε* decreases, we have increasingly lower confidence in the fact that the positives were indeed differentially expressed because of the TF perturbation. To achieve a balance between sensitivity and specificity, we evaluated three different values (i.e. 1.01, 1.05 and 1.1) for *ε*. For each TF, all ENCODE ChIP-seq peak datasets from the K562 cell line were merged to determine positives. To make the numbers of negatives and positives equal to avoid imbalanced datasets that significantly compromise the classifier performance [22], the Bray-Curtis function was applied to compute the similarity values in the expression profile between all negatives and the positive with the largest average coefficient, then the TNs with the smallest values were selected (Figure 1C).

The DHSs in the K562 cell line were intersected with known promoters. Because TFs may exhibit tissue-specific sequence preferences due to different sets of target genes and binding sites in different tissues [3], the iPWMs of EGR1, ELK1, ELF1, GABPA, IRF1, YY1 from the K562 cell line were used to most accurately detect binding sites; for ETS1, we used the only available iPWM from the GM12878 cell line [3]. Six features were derived from each homotypic cluster (i.e. Features 3 and 6 converged to the same value, because only binding sites from a single TF were used).

#### Using gene expression in the siRNA-based knockdown

In the GM19238 cell line, 59 TFs were individually knocked down using siRNAs, and significant changes in the expression levels of 8,872 genes were indicated according to their corresponding P-values [13]. In these cases, the P-value of a gene for a TF is the probability of observing the change in the expression level of this gene under the null hypothesis of no differential expression after TF knockdown; thus the larger the change in the expression level, the smaller the P-value and the more likely this gene is differentially expressed. They also indicated whether the promoters of these genes display evidence of binding to TFs by intersecting with ChIP-seq peaks in the GM12838 cell line. Among these 59 TFs, we have previously derived accurate iPWMs exhibiting primary binding motifs for 11 (BATF, JUND, NFE2L1, PAX5, POU2F2, RELA, RXRA, SP1, TCF12, USF1, YY1) [3]. Therefore, the framework for predicting TF targets in the GM19238 cell line (Figure 1A and 1D) was applied to these 11 TFs.

We defined a positive (i.e. a target gene) for a TF, if the P-value of this gene for the TF was ≤ 0.01, and the promoter interval (10kb) upstream of a TSS of this gene overlapped a ChIP-seq peak of the TF in the GM12878 cell line. A negative for a TF exhibited the following property: a P-value > 0.01 for the TF.

The DHSs in the GM19238 cell line mapped from the hg19 genome assembly were first remapped to the hg38 assembly using liftOver (available at genome.ucsc.edu) prior to being intersected with known promoters [23]. Aside from RELA, RXRA and NFE2L1, the iPWMs of TFs from the GM12878 cell line were used to detect binding sites. For RELA, the iPWM from the GM19099 cell line was used; for RXRA and NFE2L1, the only available iPWMs were respectively derived from HepG2 and K562 cells and were applied. Although the knockdown was performed in GM19238, GM12878 and GM19099 are also lymphoblastic cell lines, with GM19099 and GM19238 both being derived from Yorubans. For this analysis, the iPWMs derived in GM12878 and GM19099 were more appropriate sources of accessible TFBSs than those from HepG2 and K562, since GM12878 and GM19099 are of the same tissue type and are thus more likely comparable to GM19238 than HepG2 and K562.

### Mutation analyses on promoters of TF targets

To better understand the significance of individual binding sites for information-dense clusters and the regulatory state of direct targets, we evaluated the effects of sequence changes that altered the *R*_*i*_ values of these sites on cluster formation and whether a gene was predicted to be a TF target. Mutations were sequentially introduced into the strongest binding sites in TFBS clusters of the EGR1 target gene, *MCM7*, to determine the threshold for cluster formation after disappearing clusters disabled induction of *MCM7* expression. For one target gene of each TF from the CRISPR-generated perturbation data, effects of naturally occurring TFBS variants present in dbSNP (https://www.ncbi.nlm.nih.gov/projects/SNP/;) [24] were also evaluated to explore aspects of TFBS organization that enabled both clusters and promoter activity to be resilient to binding site mutations. This was done by analyzing whether the occurrence of individual or multiple single nucleotide polymorphisms (SNPs) lead to the loss of binding sites and the clusters that contain them, and result in changes in the predictions of these targets.

## RESULTS

### Similarity between gene expression profiles

To confirm that the Bray-Curtis Similarity can indeed effectively measure how akin the expression profiles of two genes are to one another, Equation 2 was applied to compute the similarity values between the expression profiles of the glucocorticoid receptor (*GR* or *NR3C1*) gene and all other 18,812 PC genes. *NR3C1* is an extensively characterized TF with many known direct target genes [25]. As a constitutively expressed TF activated by glucocorticoid ligands, it can mediate the up-regulation of anti-inflammatory genes by binding of homodimers to glucocorticoid response elements and down-regulation of proinflammatory genes by complexing with other activating TFs (e.g. NFKB and AP1) and eliminating their ability to bind targets [25]. NR3C1 can bind its own promoter forming an auto-regulatory loop, which also contains functional binding sites of 11 other TFs (e.g. SP1, YY1, IRF1, NFKB) whose iPWMs have been developed and/or mutual interactions have been described previously [3,25]. However, as the expression profile of *NR3C1* integrates all different splicing and translational isoforms (e.g. *GR*α*-A* to *GR*α*-D, GR*β, *GR*γ, *GR*δ), the tissue-specific expression patterns of these isoforms cannot be distinguished (e.g. levels of the *GR*α*-C* isoforms are significantly higher in the pancreas and colon, whereas levels of *GR*α*-D* are highest in spleen and lungs) [25]. *SLC25A32* and *TANK* have the greatest similarity in expression to *NR3C1 (*0.880 and 0.877 respectively), which is evident based on their overall similar expression patterns across the 53 tissues (Figure 2).

**Figure 2:**
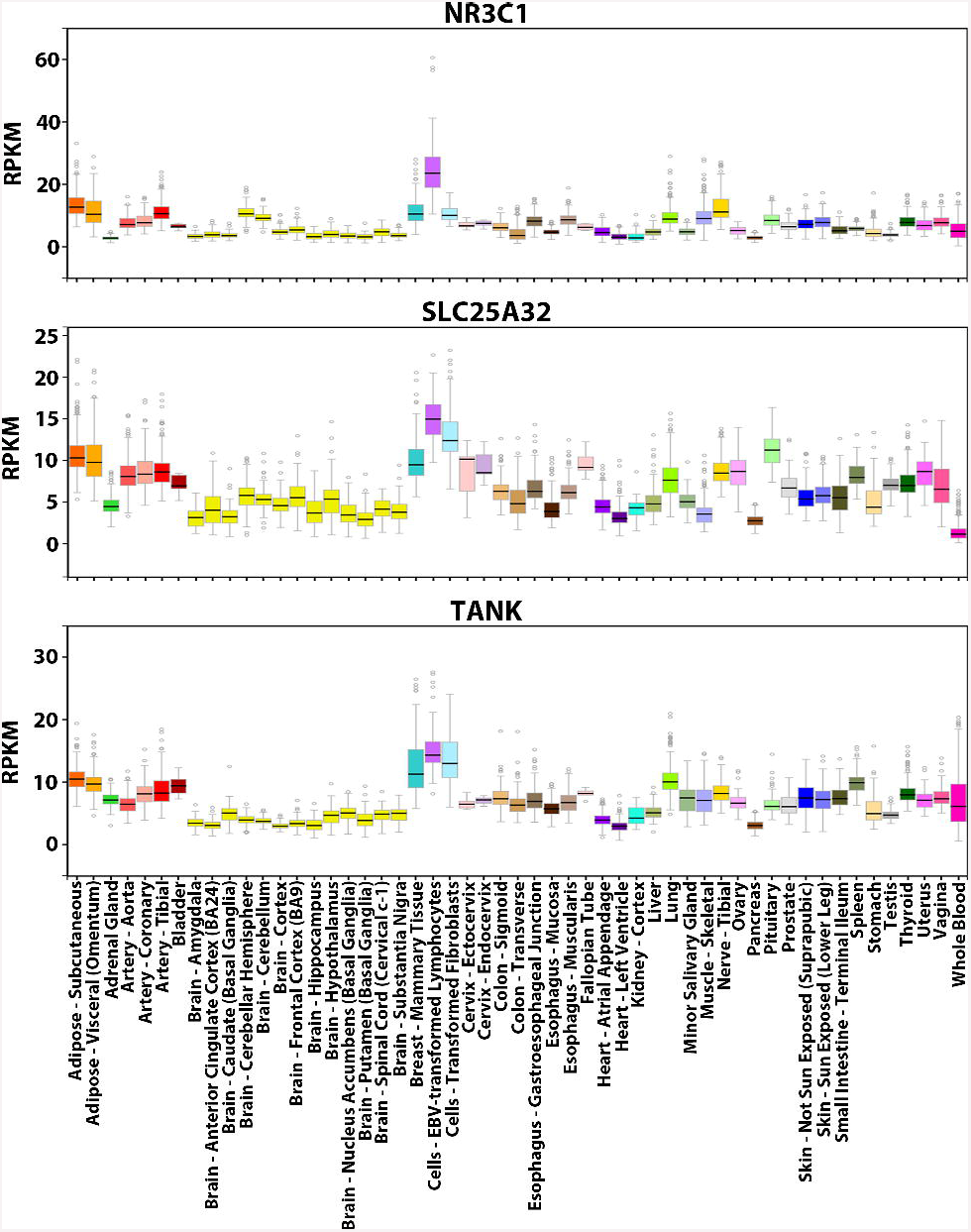
Expression profiles of NR3C1, SLC25A32 and TANK. Visualization of the expression values (in RPKM) of these genes across 53 tissues from GTEx. For each gene, the colored rectangle belonging to each tissue indicates the valid RPKM of all samples in the tissue, the black horizontal bar in the rectangle indicates the median RPKM, the hollow circles indicate the RPKM of the samples considered as outliers, and the grey vertical bar indicates the sampling error. By comparing the pictures, the overall expression patterns of the three genes across the 53 tissues resemble each other (e.g. all three genes exhibit the highest expression levels in lymphocytes and the lowest levels in brain tissues).

### Prediction of genes with similar expression profiles

In prediction of genes with similar expression profiles to *NR3C1*, we generated ROC curves to compare the performance of different classifiers (Naïve Bayes, Decision Tree, Random Forest and Support Vector Machines (SVM) with four different kernels), under two scenarios depending on whether promoter sequences were first intersected with DHSs (Figure 3). Decision Tree (DT) exhibited the largest AUC (area under the curve) under both scenarios, and was one of two most stable classifiers (i.e. ΔAUC < 0.01), with the other being the SVM with RBF kernel. Inclusion of DHS information significantly improved other classifiers’ AUC except for Naïve Bayes, and generally all TFBSs in a DHS formed a binding site cluster.

**Figure 3:**
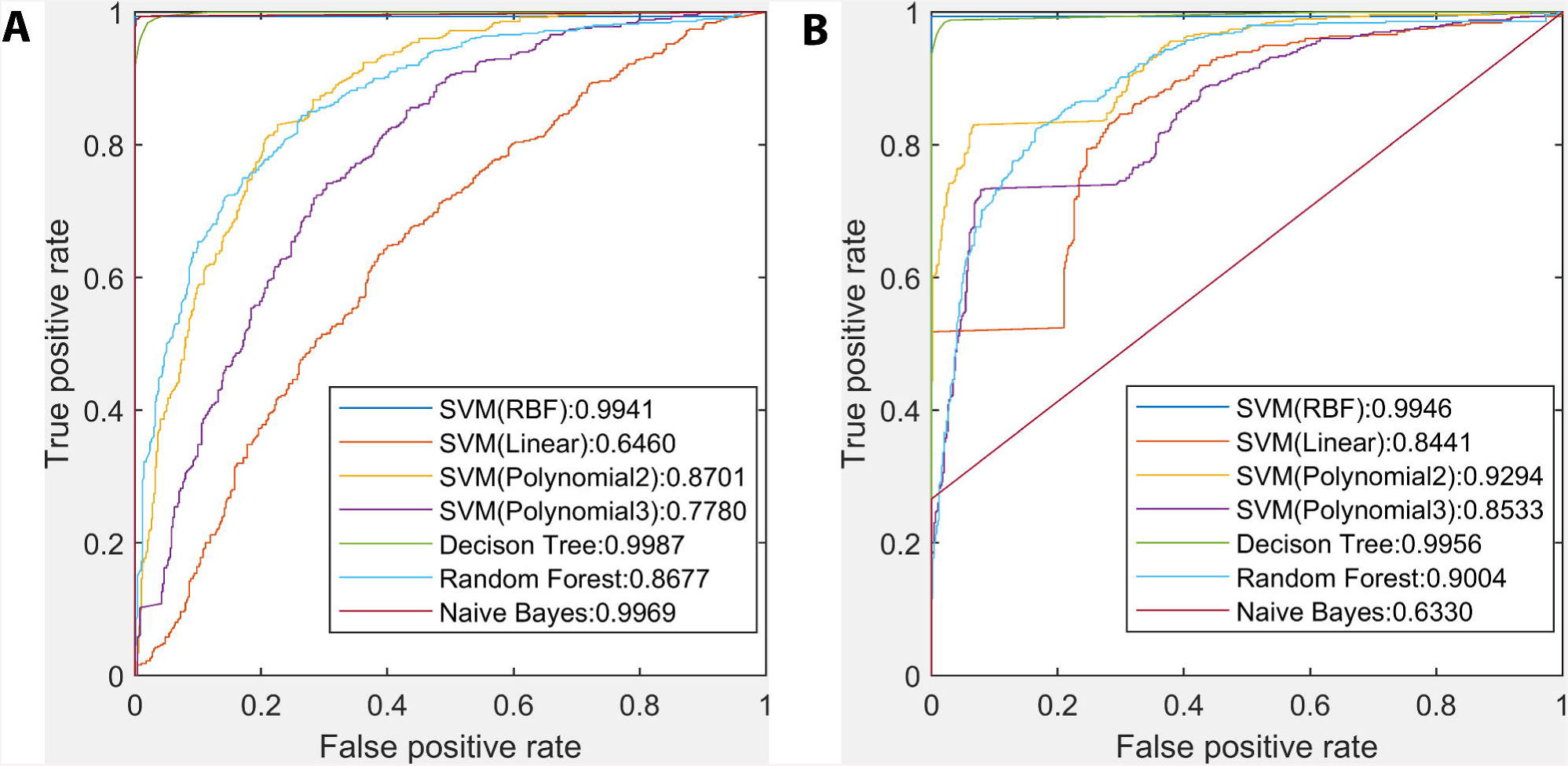
Comparison between the performance of different classifiers in prediction of genes with similar expression profiles to *NR3C1*. **A)** ROC curves and AUC of seven classifiers without intersecting promoters with DHSs. **B)** ROC curves and AUC of seven classifiers after intersecting promoters with DHSs. The Decision Tree classifier exhibited the largest AUC under both scenarios, and inclusion of DHS information significantly improved other classifiers’ AUC except for Naïve Bayes.

### Prediction of TF targets

The best-performing DT classifier in distinguishing genes with *NR3C1*-like expression profiles from others was used to predict TF targets respectively based on the CRISPR-[15] and siRNA-generated [13] perturbation data. To validate that using all six machine learning features more comprehensively capture the distribution and composition of CRMs in the promoter, feature removal was performed by only using TFBS counts. The classifier performance decreased, except for CRISPR-perturbed GABPA, IRF1 and YY1 after inclusion of DHS information (Additional file 5).

On the CRISPR-generated knockdown data, after eliminating TFBSs in inaccessible promoter intervals, i.e. those excluded from tissue-specific DHSs, the DT classifier predicted TF targets with greater sensitivity and specificity (Table 3). Specifically, predictions for TFs: EGR1, ELK1, ELF1, ETS1, GABPA, and IRF1 were more accurate than for YY1, which itself represses or activates a wide range of promoters by binding to sites overlapping the TSS (Table 3). Accordingly, the perturbation data indicated that YY1 has ∼4-22 times more PC targets in the K562 cell line than the other TFs (*ε* = 1.05), and its binding has a more significant impact on the expression levels of target genes (for YY1, the ratio of the PC target counts at *ε* = 1.1 vs *ε* = 1.01 was 0.334, which significantly exceeded those of the other TFs (0.017-0.082); Additional file 3). This is concordant with our previous finding that YY1 extensively interacts with 11 cofactors (e.g. DNA-binding IRF9 and TEAD2; non-DNA-binding DDX20 and PYGO2) in K562 cells, consistent with a central role in specifying erythroid-specific lineage development [3].

**Table 3:**
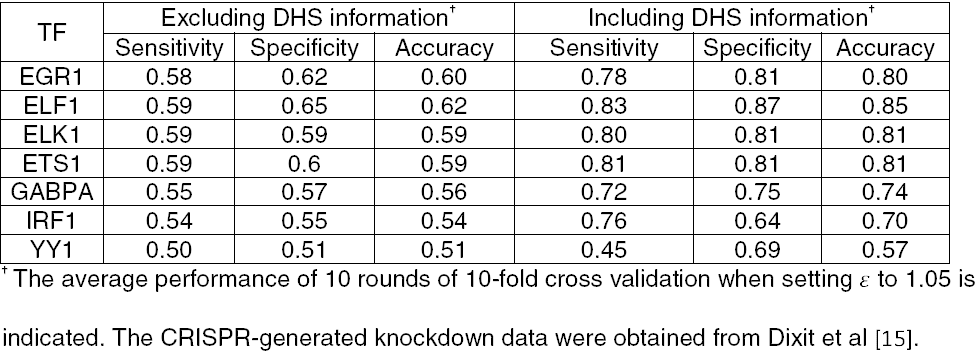
The Decision Tree classifier performance for predicting TF targets using the CRISPR-generated knockdown data.

Despite a high accuracy of target recognition, sensitivity did not exceed specificity except for IRF1 (Table 3), due to a relatively large number of false negative genes. Promoters of most TF targets contain accessible, functional binding sites that significantly change gene expression levels upon binding. By contrast, promoters of non-targets contain either no accessible binding sites at all, or accessible, but non-functional sites. The fact that these false negatives were erroneously predicted to non-targets was attributable to the indistinguishability between functional binding sites in their promoters and non-functional ones in non-targets in the classifier. *In vivo* co-regulation mediated by interacting cofactors, which was excluded by the classifier, assisted in distinguishing these non-functional sites that do not significantly affect gene expression [13].

As the threshold *ε* increased, the accuracy of the classifier for all the TFs monotonically increased as expected (Figure 4). For a gene to be defined as a DE target of a TF, the average to reach the minimum threshold *ε*. Upon TF knockdown,*ε*, is inversely correlated with the fold change in its expression level for all guide RNAs that downregulated the TF were required to reach the minimum threshold *ε*. Upon TF knockdown, *ε* is inversely correlated with the number of target genes, but positively correlated with fold changes in their corresponding expression levels. In general, more significantly DE genes have been associated with a higher number of TFBSs in their promoters [13]. Thus, at greater *ε*, there are larger differences in the values of machine learning features derived from TFBS clusters between direct targets and non-targets.

**Figure 4:**
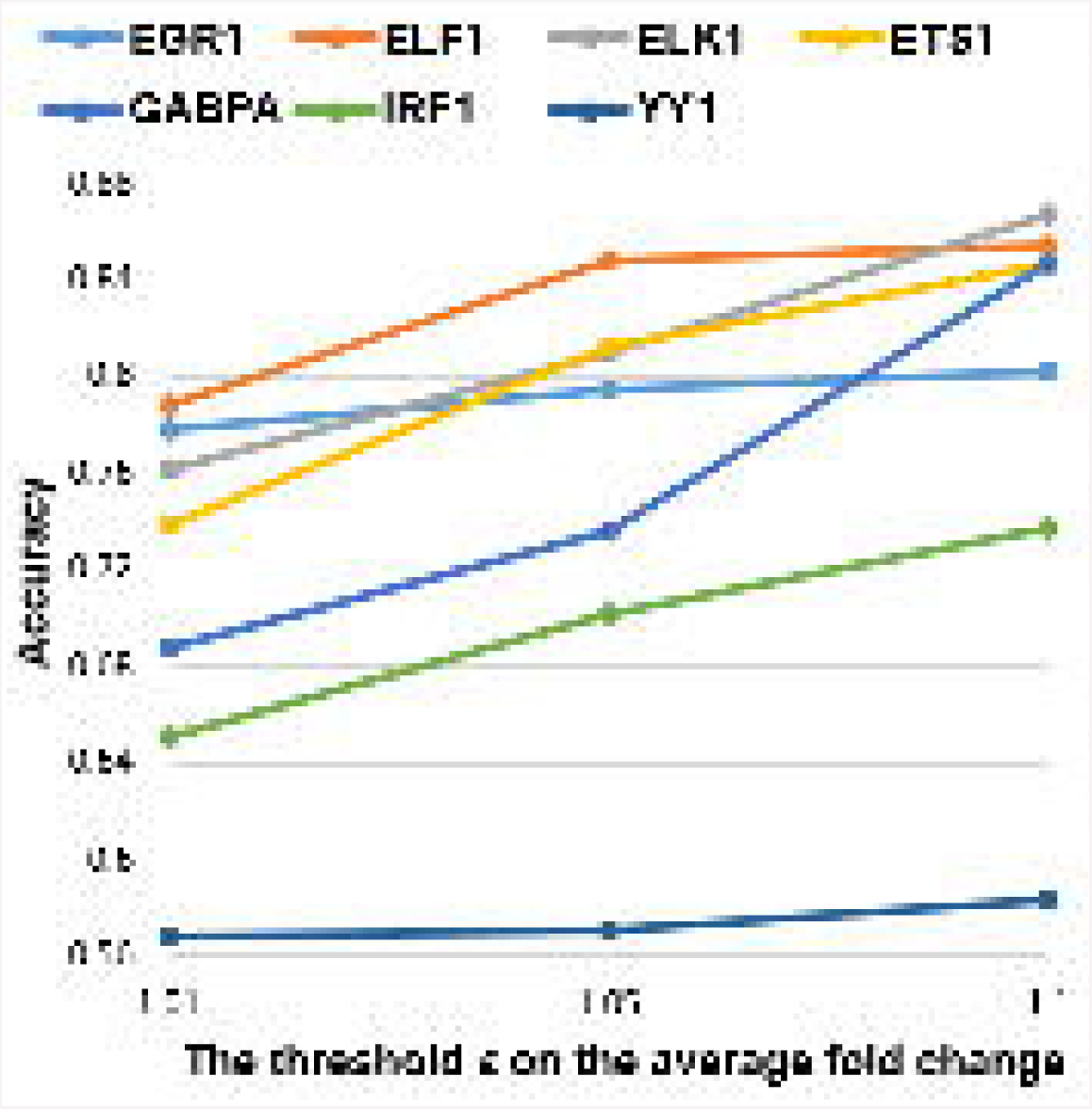
Accuracy of the Decision Tree classifier when using three different values for *ε*. Each accuracy value was averaged from 10 rounds of 10-fold cross validation, when the minimum threshold, on the average fold change in gene expression levels under all guide Each accuracy value was averaged from 10 rounds of 10-fold cross validation, when the RNAs of the TF took three different values 1.01, 1.05 and 1.1. As, increased, accuracy for all seven TFs monotonically increased.

With the siRNA-generated knockdown data, the performance of the DT classifier was compared to the approach inferring DE targets by correlating TF binding with gene expression levels across ten cell types [14]. In this correlation-based approach, three measures (i.e. the absolute Pearson correlation coefficient (PCC), the absolute Spearman correlation coefficient (SCC), and the absolute combined angle ratio statistic (CARS)), whose performance was evaluated with precision-recall curves, were alternatively used to compute a correlation score between the number of ChIP-seq peaks overlapping the promoter and gene expression values. Genes predicted to be DE targets had scores above the threshold resulting in a 1.5-fold increase compared to the background precision (i.e. the DE target count / 8,872). For example, in the case of YY1, which was the only TF analyzed by both approaches, the performance of the DT classifier was 0.98 (precision) and 0.55 (recall) after including DHS information (Table 4). This classifier outperformed all three correlation measures (PCC: 0.467 and 0.003; SCC: 0.467 and 0.006; CARS: 0.467 and 0.003 directly obtained from [14]), even though the correlation-based approach used a less stringent P-value threshold (0.05) for defining differential expression of likely non-direct targets, and intersected ChIP-seq peaks over shorter 5kb promoter intervals upstream of the TSS. Three reasons explain why the correlation-based approach exhibited lower recall, including that 1) it did not use machine learning classifiers, 2) its larger P-value threshold (0.05) generated a larger number of positives and 3) its positives also include DE targets that cannot be directly bound.

**Table 4:** The Decision Tree classifier performance for predicting TF targets using the siRNA-generated knockdown data.

| TF | Excluding DHS information<sup>†</sup> |  |  | Including DHS information<sup>†</sup> |  |  |
| --- | --- | --- | --- | --- | --- | --- |
|  | Sensitivity | Specificity | Accuracy | Sensitivity | Specificity | Accuracy |
| BATF | 0.96 | 0.97 | 0.96 | 0.85 | 1 | 0.93 |
| JUND | 0.86 | 0.90 | 0.88 | 0.80 | 1 | 0.90 |
| NFE2L1 | 0.92 | 0.95 | 0.94 | 0.71 | 0.93 | 0.82 |
| PAX5 | 0.96 | 0.97 | 0.96 | 0.88 | 0.98 | 0.93 |
| POU2F2 | 0.97 | 0.97 | 0.97 | 0.89 | 0.99 | 0.94 |
| RELA | 0.95 | 0.96 | 0.96 | 0.83 | 0.97 | 0.90 |
| RXRA | 0.93 | 0.91 | 0.92 | 0.84 | 0.95 | 0.89 |
| SP1 | 0.98 | 0.98 | 0.98 | 0.89 | 0.99 | 0.94 |
| TCF12 | 0.98 | 0.98 | 0.98 | 0.86 | 0.99 | 0.93 |
| USF1 | 0.97 | 0.98 | 0.97 | 0.83 | 0.98 | 0.90 |
| YY1 | 1 | 1 | 1 | 0.55 | 0.99 | 0.77 |

<sup>†</sup> The average performance of 10 rounds of 10-fold cross validation is indicated. The siRNA-generated knockdown data were obtained from Cusanovich et al [13].

### Intersection of genes with similar expression profiles and TF targets

To determine how many TF targets have similar tissue-wide expression profiles, we intersected the set of targets with the set of 500 PC genes with the most similar expression profiles for each TF (Table 5, Additional file 6). The TFs PAX5 and POU2F2 are primarily expressed in B cells, and their respective targets *IL21R* and *CD86* are also B cell-specific, which accounts for the high similarity in the expression profile between them. There are respectively 21 and 7 nuclear mitochondrial genes (e.g. *MRPL9* and *MRPS10*, which are subunits of mitochondrial ribosomes) in the intersections for YY1 in the K562 and GM19238 cell lines [26]. Previous studies reported that YY1 upregulates a large number of mitochondrial genes by complexing with PGC-1α in C2C12 cells [27], and genes involved in the mitochondrial respiratory chain in K562 cells [15], which is consistent with the idea that YY1 may broadly regulate mitochondrial function (within all 53 tissues in addition to the erythrocyte, lymphocyte and skeletal muscle cell lines).

**Table 5:** Intersection of TF targets and 500 protein-coding genes with the most similar expression profiles.

| TF | Cell line | Number of targets | Size of intersection | Targets among the most similar 10 genes<sup>§</sup> |
| --- | --- | --- | --- | --- |
| EGR1 | K562 | 169 | 12 | None |
| ELF1 |  | 78 | 5 | None |
| ELK1 |  | 112 | 4 | GNL1(8<sup>th</sup>) |
| ETS1 |  | 267 | 15 | None |
| GABPA |  | 513 | 25 | TAF1(1<sup>st</sup>) |
| IRF1 |  | 457 | 10 | None |
| YY1 |  | 1752 | 127 | MRPL9(2<sup>nd</sup>), BAZ1B(6<sup>th</sup>), ENY2(7<sup>th</sup>),<br>NUB1(8<sup>th</sup>), USP1(9<sup>th</sup>), HNRNPR(10<sup>th</sup>) |
|  | GM19238 | 1040 | 61 | MED4(1<sup>st</sup>), SURF6(3<sup>rd</sup>), BAZ1B(6<sup>th</sup>) |
| BATF |  | 186 | 21 | None |
| JUND |  | 44 | 2 | None |
| NFE2L1 |  | 58 | 4 | None |
| RELA |  | 247 | 13 | HMG20B(9<sup>th</sup>) |
| RXRA |  | 181 | 3 | None |
| SP1 |  | 1595 | 81 | None |
| TCF12 |  | 655 | 20 | None |
| USF1 |  | 301 | 21 | None |
| PAX5 |  | 918 | 86 | IL21R(8<sup>th</sup>) |
| POU2F2 |  | 532 | 26 | CD86(3<sup>rd</sup>) |

<sup>§</sup> The rank of each target in the list of similar genes in the descending order of Bray-Curtis similarity values is shown in the brackets immediately following the target.

Between 0.4%-25% of genes with similar expression profiles to the TFs are actually their targets (Table 5); the majority are non-targets whose promoters contain non-functional binding sites that are distinguished from targets by their lack of coregulation by corresponding cofactors. For YY1 and EGR1, we validated this hypothesis by contrasting the flanking cofactor binding site distributions and strengths in the promoters of the most similarly expressed target genes (YY1: *MRPL9, BAZ1B*; EGR1: *CANX, NPM1*) and non-target genes (YY1: *ADNP, RNF25*; EGR1: *GUCY2F, AWAT1*). Strong and intermediate recognition sites for TFs: SP1, KLF1, CEBPB formed heterotypic clusters with adjacent YY1 sites; as well TFBSs of SP1, KLF1, and NFY were frequently present adjacent to EGR1 binding sites. These patterns contrasted with the enrichment of CTCF and ETV6 binding sites in gene promoters of YY1 and EGR1 non-targets (Additional file 7). Previous studies have reported that KLF1 is essential for terminal erythroid differentiation and maturation [26], direct physical interactions between YY1 and the constitutive activator SP1 synergistically induce transcription [29], the activating CEBPB promotes differentiation and suppresses proliferation of K562 cells by binding the promoter of the *G-CSFR* gene encoding a hematopoietin receptor [30], EGR1 and SP1 synergistically cooperate at adjacent non-overlapping sites on *EGR1* promoter but compete binding at overlapping sites [31]; whereas CTCF binding sites function as an insulator blocking the effects of *cis*-acting elements and preventing gene activation by mediating long-range DNA loops to alter topological chromatin structure [32,33], and ETV6, a member of the ETS family, is a transcriptional repressor required for bone marrow hematopoiesis and associated with leukemia development [34].

### Mutation analyses on promoters of direct targets

Because the promoters of most direct targets contain multiple binding site clusters, we anticipate that this enables these genes’ expression to be naturally robust against binding site mutations; in other words, the other clusters can compensate for the loss of a cluster destroyed by mutations in binding sites, so that the mutated promoters are still capable of effectively inducing gene transcription upon TF binding. First, we validated this hypothesis *in silico* by examining whether introducing artificial variants into binding sites in the promoter of the target gene *MCM7* of EGR1 changes the classifier output (Figure 5). Specifically, in the K562 cell line, *MCM7* is upregulated by EGR1. Knockdown of *MCM7* has an anti-proliferative and pro-apoptotic effect on K562 cells [35] and the loss of EGR1 increases leukemia initiating cells [36], which suggests that EGR1 may act as a tumor suppressor in K562 cells through the *MCM7* pathway.

**Figure 5:**
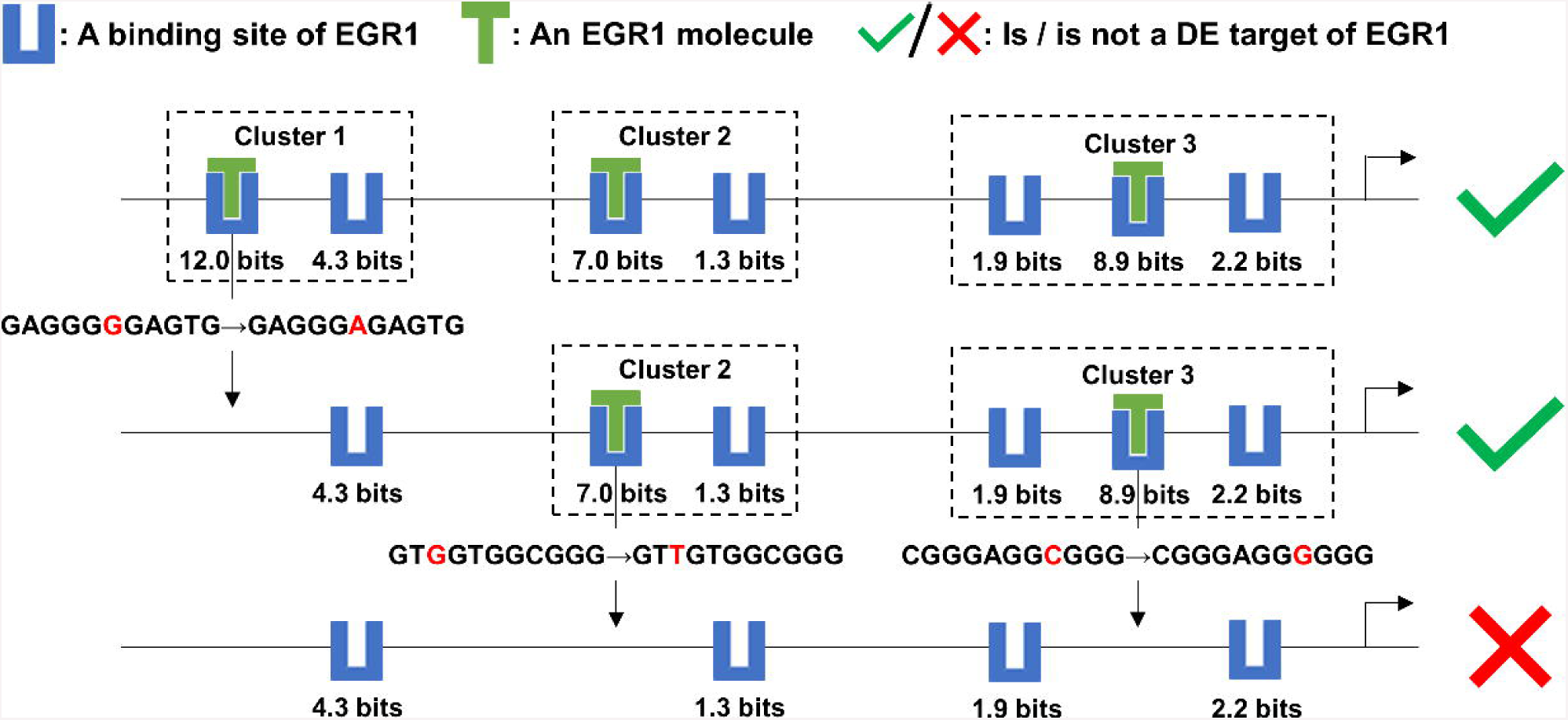
Mutation analyses on the target MCM7 of EGR1. This figure depicts the effect of a mutation in each EGR1 binding site cluster of the MCM7 promoter on the expression level of MCM7, which is a target of the TF EGR1. The strongest binding site in each cluster were abolished by a single nucleotide variant. Upon loss of all three clusters, only weak binding sites remained and EGR1 was predicted to no longer be able to effectively regulate MCM7 expression. Multiple clusters in the promoters of TF targets confer robustness against mutations within individual binding sites that define these clusters.

First, the strongest binding site (at position chr7:100103347 [hg38], -strand, *R*_*i*_ = 12.0 bits) in the promoter was eliminated by a G->A mutation, resulting in the disappearance of Cluster 1, which consists of two sites (the other site at chr7:100103339, -, 4.3 bits). EGR1 was still predicted to compensate for this mutation, due to the presence of the other two clusters comprising weaker binding sites of intermediate strength (chr7:100102252, +, 7.0 bits; chr7:100102244, +, 1.3 bits; chr7:100101980, +, 8.9 bits; chr7:100101977, +, 2.2 bits; chr7:100101984, +, 1.9 bits), enabling the promoter to maintain capability of inducing *MCM7* expression (Figure 5). These adjacent clustered sites, which may not be strong enough to bind TFs and individually activate transcription, can stabilize each other’s binding [2]. The weaker sites flanking a strong binding site in a cluster can direct the TF molecule to the strong site and extend the period of the molecule physically associating with the strong site, which is termed, the funnel effect [2]. Further, Clusters 2 and Cluster 3 were respectively removed by G->T and C->G mutations abolishing the strongest site in either cluster, which altered the prediction, that is, EGR1 lost the capability to induce *MCM7* transcription (Figure 5). The remaining four sparse weak sites do not form a cluster and cannot completely supplant the disrupted strong sites.

Further, we examined the impacts of known natural SNPs on binding site strengths, clusters and the regulatory state of the promoter for a direct target of each of the seven TFs from the CRISPR-generated perturbation data (Table 6). Often a single SNP (e.g. rs996639427 of EGR1) can affect the strengths of multiple binding sites (Table 6). Apart from SNPs that are predicted to abolish binding (Figure 5), leaky variants that merely weaken TF binding are common (Table 6). Binding stabilization between adjacent sites and the funnel effect enable the CRMs comprised of information-dense clusters to be robust to mutations in individual binding sites. In this way, neither mutations that abolish TFBSs nor leaky SNPs in flanking weak sites can destroy functional clusters (e.g. rs1030185383 and rs5874306 of IRF1), whereas SNPs with large reductions in *R*_*i*_ values of central strong sites are more likely to abolish clusters (e.g. rs865922947, rs946037930, rs917218063 and rs928017336 of YY1) (Table 6). More generally, the presence of multiple clusters enables promoters to be effectively resilient to the effects binding site mutations; only the complete abolishment of all clusters resulting from the simultaneous occurrence of multiple SNPs can transform the promoter to be unresponsive to TF binding to residual weak sites (e.g. rs997328042, rs1020720126 and rs185306857 of GABPA) (Table 6). Furthermore, a relatively small number of SNPs that strengthen TF binding and eventually amplify the regulatory effect of the TF on the gene expression level are also present (e.g. rs887888062 of EGR1 and rs751263172 of ELF1) (Table 6), suggesting that, in addition to deleterious mutations, benign variants may also be found in promoters, consistent with the expectations of neutral theory [37].

**Table 6:**
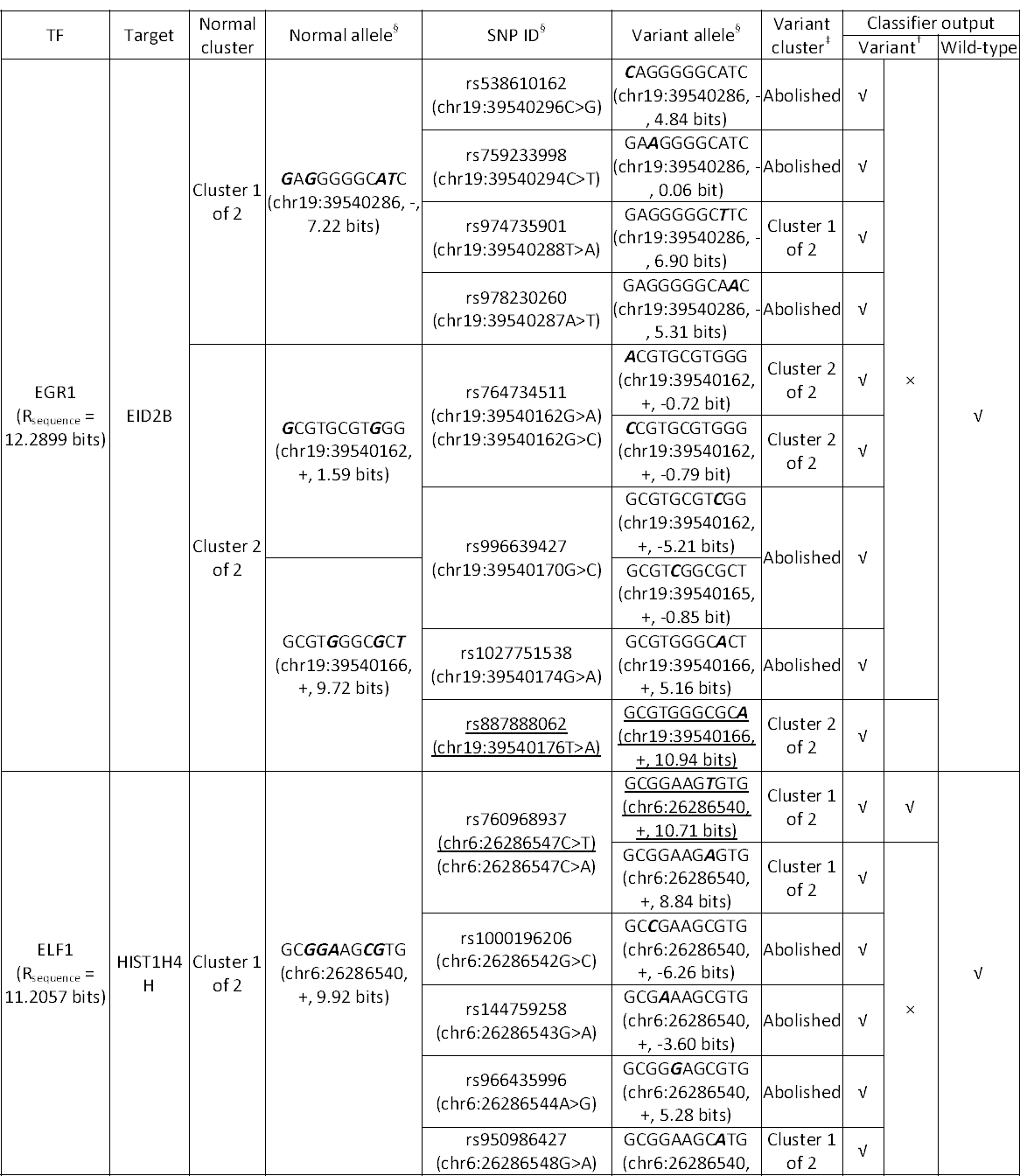
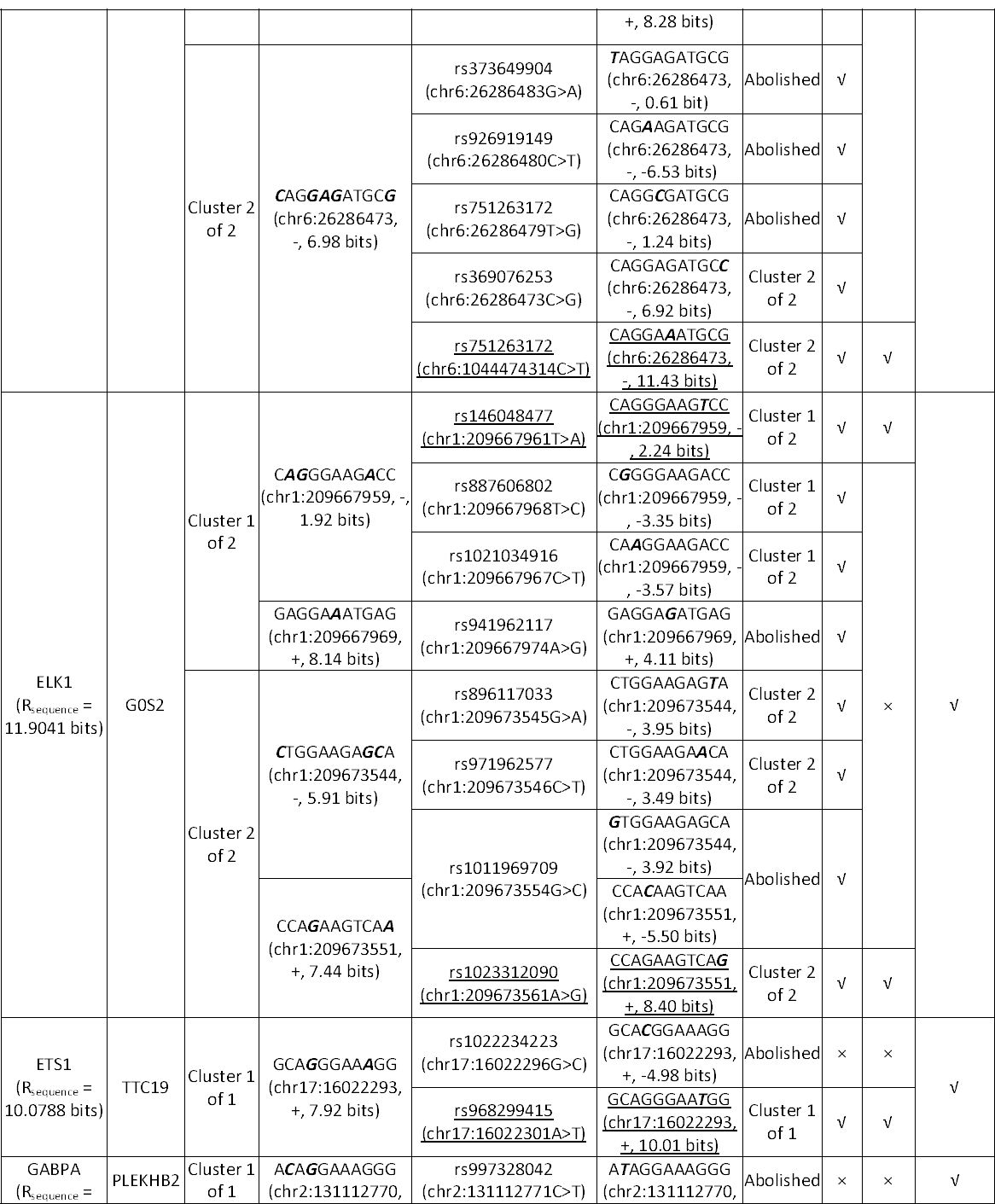
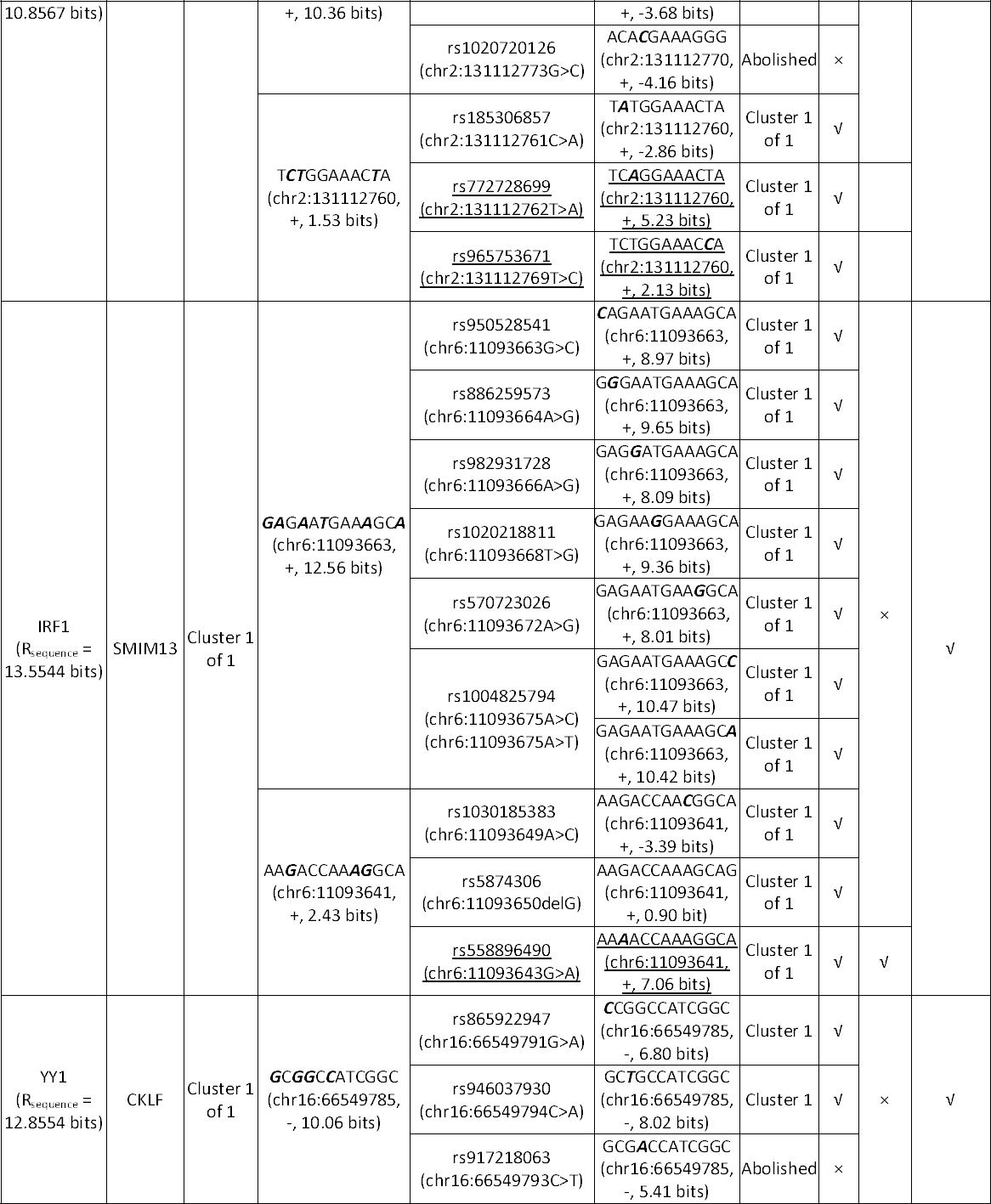
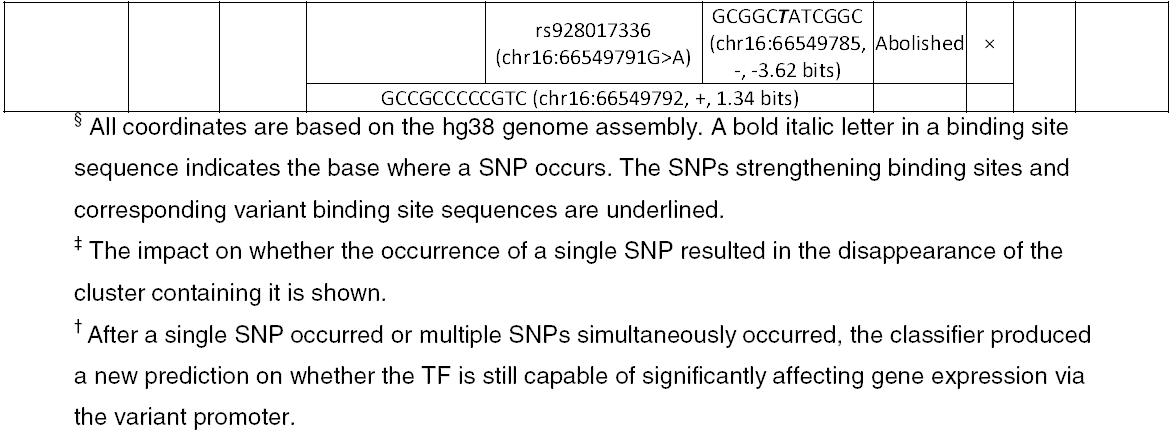
Mutation analyses on promoters of TF targets.

## DISCUSSION

In this study, the Bray-Curtis similarity function was initially shown (for the *NR3C1* gene) to measure the relatedness of overall expression patterns between genes across a diverse set of tissues. The resulting machine learning framework distinguished Bray-Curtis function-defined similar from dissimilar genes based on the distribution, strengths and compositions of TFBS clusters in accessible promoters, which can substantially account for the corresponding gene expression patterns. Using knockdown data as the gold standard, the combinatorial use of TF binding profiles and chromatin accessibility was also demonstrated to be predictive of TF targets. A binding site comparison confirmed that coregulatory cofactors can be used to responsible for distinguishing between functional sites in targets and non-functional ones in non-targets. Furthermore, mutation analyses on binding sites of targets demonstrated that the existence of both multiple TFBSs in a cluster and multiple information-dense clusters in a promoter enables both the cluster and the promoter to be resilient to binding site mutations.

The DT classifier improved after intersecting promoters with DHSs in prediction of TF targets with the CRISPR-generated knockdown data (Table 3). This intersection eliminated noisy binding sites that are inaccessible to TF proteins in promoters; specifically, it widened discrepancies in feature vectors between positives and negatives. If the 10kb promoter of a gene instance does not overlap DHSs, its feature vector will only consist of 0; the percentages of negatives whose promoters do not overlap DHSs considerably exceeded those of positives (Additional file 8), which led to an excess of negatives with feature vectors containing only 0 after intersection. This explains why these negatives are not DE targets of the TFs in the K562 and GM19238 cell lines, because their entire promoters are not open to TF molecules; other regulatory regions besides the proximal promoters (e.g. intronic enhancers [38]) still enable the TFs to effectively control the expression of the positives with inaccessible promoters. The relatively poor performance of the classifier on YY1 (Table 3) is attributable to its smaller percentage of negatives with inaccessible promoters (Additional file 8) and the larger number of functional binding sites in the K562 cell line.

Our *in-silico* mutation analyses revealed that some deleterious TFBS mutations could be compensated for by other information-dense clusters in a promoter; thus predicting the effects of mutations in individual binding sites would not be sufficient to interpretation of downstream effects. Though compensatory clusters may maintain gene expression, the promoter will provide lower levels of activity than the wild-type promoter could, which is a recipe for achieving natural phenotypic diversity. Few published studies in molecular diagnostics have specifically examined the effects of naturally occurring variants within clustered TFBSs; thus IDBC-based machine learning provided an alternative computational approach to predict deleterious mutations that actually impact (i.e. repress or abolish) transcription of target genes and result in abnormal phenotypes, and to simultaneously minimize false positive calls of TFBS mutations that individually have little or no impact.

Apart from these TFs, the Bray-Curtis Similarity can be directly applied to identify the ground-truth genes with overall similar tissue-wide expression patterns to any other gene whose expression profile is known. Further studies could investigate the biological significance underlying the phenomenon that all these genes share a common expression pattern, including the similarity between other regulatory regions besides proximal promoters in terms of TFBSs and epigenetic markers. This machine learning framework can also be applied to predict target genes for other TFs and in other cell lines, depending on the availability of corresponding knockdown data.

There are a number of limitations of our approach. The Bray-Curtis function seems unable to accurately measure the similarity between the expression profiles of a gene (e.g. *MIR23A*) without any detectable mRNA in any of the 53 tissues analyzed and genes (e.g. the ubiquitously expressed *NR3C1* and stomach-specific *PGA3*) that are expressed in at least one tissue. Intuitively, in terms of expression patterns *PGA3* is more similar to *MIR23A* than *NR3C1*; however, the Bray-Curtis similarity values indicate that both *PGA3* and *NR3C1* bear no similarity to *MIR23A (*i.e. *sim*_Bray-Curtis_ (*NR*3*C*1,*MIR*23*A*)=*sim*_Bray-curtis_ (*PGA*3, *MIR*23*A*) = 0). Another possible limitation in classifier performance in the prediction of genes with similar tissue-wide expression profiles is that only binding sites of 82 TFs were analyzed due to a lack of available iPWMs for other TFs, given that 2000-3000 sequence-specific DNA-binding TFs are estimated to be encoded in the human genome [39]. For example, four TFs (CREB, MYB, NF1, GRF1) were previously reported to bind the promoter of the *NR3C1* gene to activate or repress its expression, however their iPWMs exhibiting known primary motifs could not be successfully derived from ChIP-seq data [3,25]. Regarding the CRISPR-generated knockdown data used here, positives were inferred to be direct targets by intersecting their promoters with corresponding ChIP-seq peaks, which may not be completely accurate, due to the presence of noise peaks that do not contain true TFBSs [3,40]. Small fold changes in the expression levels of DE targets could arise from compromised efficiency of knockdowns as a result of suboptimal guide RNAs or the limitations of perturbing only a single allele of the TF. Finally, the framework developed here only takes into account the 10kb interval proximal to the TSS, and would not therefore capture long range enhancer effects beyond this distance; by contrast, correlation based approaches have successfully incorporated multiple definitions of promoter length [14].

## CONCLUSIONS

The Bray-Curtis function is able to effectively quantify the similarity between tissue-wide gene expression profiles. By analysis of information theory-based TF binding profiles that captured the spatial distribution and information contents of TFBS clusters, ChIP-seq and chromatin accessibility data, we described a machine learning framework that distinguished tissue-wide expression profiles of similar vs dissimilar genes and identified TF target genes. Functional binding sites in target genes that significantly alter expression levels upon direct binding are at least partially distinguished by TF-cofactor coregulation from non-functional sites in non-targets. Finally, *in-silico* mutation analyses demonstrated that the presence of multiple information-dense clusters in the promoter, as a protective mechanism, reduces deleterious mutations that can significantly alter the regulatory state and expression level of the gene.

## LIST OF ABBREVIATIONS

TF: transcription factor
TFBS: transcription factor binding site
CRM: *cis*-regulatory modules
iPWM: information theory-based position weight matrix
IDBC: information density-based clustering
*R*_*i*_: Rate of Shannon information transmission for an individual sequence
ChIP-seq: chromatin immunoprecipitation with massively parallel DNA sequencing
HM: histone modification
mRNA: messenger RNA
siRNA: small interfering RNA
CRISPR: clustered regularly interspaced short palindromic repeats
DE: differentially expressed
DHS: deoxyribonuclease I hypersensitive region
RPKM: reads per kilobase of transcript per million mapped reads
GR: glucocorticoid receptor
GTEx: genotype-tissue expression
ENCODE: encyclopedia of DNA elements
TSS: transcription start site
SVM: support vector machine
RBF: radial basis function
DT: decision tree
ROC: receiver operating characteristic
AUC: area under the curve
PC: protein-coding
PCC: absolute Pearson correlation coefficient
SCC: absolute Spearman correlation coefficient
CARS: the absolute combined angle ratio statistic
SNP: single nucleotide polymorphism.

## ADDITIONAL FILES

### Additional file 1

The mathematical definitions of the four other similarity metrics, the workflow of the IDBC algorithm, the mathematical definitions of five statistical variables to measure classifier performance, and the default parameter values of classifiers in MATLAB

Format:.docx

### Additional file 2

The lists of positives and negatives in the machine learning classifiers to predict genes with similar tissue-wide expression profiles

Format:.xlsx

### Additional file 3

The lists of positives and negatives in the DT classifier to predict TF targets based on the CRISPR-generated knockdown data

Format:.xlsx

### Additional file 4

The lists of positives and negatives in the DT classifier to predict DE direct targets based on the siRNA-generated knockdown data

Format:.xlsx

### Additional file 5

The performance of the DT classifier using only TFBS counts Format:.xlsx

### **Additional file 6**

The list of the most similar 500 PC genes to each TF in terms of expression profiles, and the intersection of these 500 genes and target genes of the TF

Format:.xlsx

### **Additional file 7**

Cofactor binding sites adjacent to YY1 and EGR1 sites in the promoters of their targets and non-targets

Format:.docx

### Additional file 8

The percentages of positives and negatives whose promoters do not overlap DHSs for the CRISPR-perturbed TFs

Format:.xlsx

## DECLARATIONS

### Ethics approval and consent to participate

Not applicable

### Consent for publication

Not applicable

### Availability of data and materials

The median RPKM, TSS coordinate, DNase I hypersensitivity and ChIP-seq data are respectively available from the GTEx Analysis V6p release (www.gtexportal.org), Ensembl Biomart (www.ensembl.org) and ENCODE (www.encodeproject.org). The CRISPR- and siRNA-generated knockdown data are available from the supplementary information files of Dixit et al. [15] and Cusanovich et al. [13]. The code implementing this machine learning framework and the datasets generated and/or analysed by this framework are available in Zenodo (https://doi.org/10.5281/zenodo.1249635). All other data generated or analysed during this study are included in this published article and its supplementary information files.

## Authors’ contributions

PKR defined the objectives and directed the study. RL and PKR devised the general machine learning framework. RL implemented this framework and collected the results. Both RL and PKR interpreted the results and wrote the manuscript.

## Acknowledgements

We are grateful to Ben Shirley and Eliseos Mucaki for constructive comments on the paper.

