## Supplementary Materials for "Information-dense transcription factor binding site clusters identify target genes with similar tissue-wide expression profiles and serve as a buffer against mutations"

**1) Mathematical definitions of the four other similarity metrics in Table 1**

Euclidean Similarity:

$${sim}_{Euclidean}\left( {EP}^{A},{EP}^{B} \right)=\frac{1}{1+d_{Euclidean}\left( {EP}^{A},{EP}^{B} \right)}=\frac{1}{1+\sqrt{\sum_{i=1}^{53} \left( {MEV}_{i}^{A}-{MEV}_{i}^{B} \right)^{2}}}$$

Cosine Similarity:

$${sim}_{Cosine}\left( {EP}^{A},{EP}^{B} \right)=\frac{\sum_{i=1}^{53} {MEV}_{i}^{A}{MEV}_{i}^{B}}{\sqrt{\sum_{i=1}^{53} \left( {MEV}_{i}^{A} \right)^{2}}\sqrt{\sum_{i=1}^{53} \left( {MEV}_{i}^{B} \right)^{2}}}$$

Pearson correlation:

$${sim}_{Pearson}\left( {EP}^{A},{EP}^{B} \right)=\frac{\sum_{i=1}^{53} \left( {MEV}_{i}^{A}-\bar{{MEV}^{A}} \right)\left( {MEV}_{i}^{B}-\bar{{MEV}^{B}} \right)}{\sqrt{\sum_{i=1}^{53} \left( {MEV}_{i}^{A}-\bar{{MEV}^{A}} \right)^{2}}\sqrt{\sum_{i=1}^{53} \left( {MEV}_{i}^{B}-\bar{{MEV}^{B}} \right)^{2}}}$$

where ${EP}^{A}$ and ${EP}^{B}$ are respectively the expression profiles of Gene $A$ and $B$ (Equation 1).

Spearman correlation:

First the median expression values in ${EP}^{A}$ and ${EP}^{B}$ are ranked separately. In each expression profile, the highest expression value is labelled "1" and the lowest value is labelled "53". In the case that there are multiple identical values, each value will take the average of the ranks that they would have otherwise occupied.

$${sim}_{Spearman}\left( {EP}^{A},{EP}^{B} \right)=1-\frac{6\sum_{i=1}^{53} d_{i}^{2}}{53\left( {53}^{2}-1 \right)}$$

where $d_{i}$ is the difference in the paired ranks corresponding to Tissue $i$.

**2) The Information density-based clustering (IDBC) algorithm** [1]

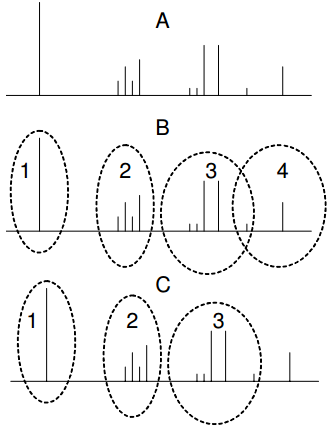

**Figure 1.** An example of the IDBC algorithm running on a promoter**.** Panel A shows the locations of putative binding sites upstream of the transcription start site (TSS). The vertical height of each bar indicates the strength of the respective binding site. Panel B shows the initial list of 4 clusters derived from the first iteration of the algorithm. This includes an example of an overlap where one of the sites is shared between Clusters 3 and 4. Panel C shows the result of a refining step where the overlapping site is resolved by putting it into Cluster 3. Since the single site in Cluster 4 is not strong enough to be a cluster, finally only 3 clusters remain.

The detailed steps of this algorithm are:

1. For each TFBS *s*, calculate the neighborhood information content (*nic*) as being the total of pairwise sums of the information content for *s* and each site lying within a distance *d* (number of bases) of *s*.

2. For every TFBS *s* that has an *nic* exceeding a threshold parameter *I*, create an initial cluster by promoting s to the role of the cluster center *c* and including all TFBSs within its neighborhood as members. (Figure 1)

3. In the first phase of merging clusters (Figure 1), consider all pairs of clusters with centers *c_i_* and *c_j_*.

- If *c_i_* is a member of the cluster with *c_j_* as its center and vice versa, then merge the two clusters and replace the center with the stronger one of *c_j_* and *c_j_*. If they are equal in strength, the center of the cluster containing more TFBSs is made the center of the new cluster, while the other center is relegated to being just a site.
- If *c_i_* is a member of the cluster with *c_j_* as its center and *c_j_* is not a member of the cluster with *c_i_* as its center, compare the strengths of *c_i_* and *c_j_*.
  - If *c_i_* is stronger than *c_j_*, the overlapping TFBSs are put into the cluster with *c_i_* as its center and removed from the cluster with *c_j_* as its center. Among the remaining TFBSs in the cluster with *c_j_* as its center, the strongest TFBS *c_k_* is selected as the new center.
  - If *c_i_* is weaker than *c_j_*, the overlapping TFBSs are put into the cluster with *c_j_* as its center and removed from the cluster with *c_i_* as its center.

This process is iterated until no *c_i_* occurs in more than a single cluster.

4. In the second phase of merging (Figure 1), all TFBSs that belong to more than one cluster are exclusively allocated to the cluster with the stronger center.

5. In the re-evaluation phase (Figure 1), a final check is made to ensure that each cluster fulfils the criterion of minimum information density *I* (as in step 2) after the possible reallocation of sites in the preceding step. Clusters failing the check are dissolved into individual sites.

**3) Mathematical definitions of the statistical variables to measure classifier performance**

Table 1. The confusion matrix output by a machine learning classifier on a training/test set

| **Total population** | | **True condition** | |
| --- | --- | --- | --- |
|  |  | Condition positive | Condition negative |
| **Predicted**  **condition** | Predicted condition positive | True positive (TP) | False positive (FP) |
|  | Predicted condition negative | False negative (FN) | True negative (TN) |

The formal definitions of five statistical variables are given below:

Accuracy = $\frac{TP + TN}{Total population}$ = $\frac{TP + TN}{TP + TN + FP + FN}$

Sensitivity = Recall = $\frac{TP}{Condition positive}$ = $\frac{TP}{TP + FN}$

Specificity = $\frac{TN}{Condition negative}$= $\frac{TN}{TN + FP}$

Precision = $\frac{TP}{Predicted condition positive}$ = $\frac{TP}{TP + FP}$

**4) Default parameters of the seven classifiers in MATLAB 2018a**

① SVMs:

| Parameter | Default value |  | Parameter | Default value |
| --- | --- | --- | --- | --- |
| Box constraint | 1 |  | Standardize | false |
| Kernel scale | 1 |  | Solver | ISDA |
| Kernel offset | 0 |  | Alpha | zeros(*n*,1) |
| Cache size | 1000 |  | Clip alphas | true |
| NumPrint | 1000 |  | Outlier fraction | 0.0 |
| Remove duplicates | false |  |  |  |

where *n* is the number of instances.

Kernel functions:

RBF kernel: $G\left( x_{j},x_{k} \right)=exp\left( -\left\| x_{j}-x_{k} \right\|^{2} \right)$

Linear kernel: $G\left( x_{j},x_{k} \right)=x_{j}^{'}x_{k}$

Polynomial kernel of order 2: $G\left( x_{j},x_{k} \right)=\left( 1+x_{j}^{'}x_{k} \right)^{2}$

Polynomial kernel of order 3: $G\left( x_{j},x_{k} \right)=\left( 1+x_{j}^{'}x_{k} \right)^{3}$

② Decision Tree:

| Parameter | Default value |  | Parameter | Default value |
| --- | --- | --- | --- | --- |
| Maximum depth | +∞ |  | MinParentSize | 10 |
| MergeLeaves | on |  | PredictorSelection | allsplits |
| Prior | empirical |  | Prune | off |
| PruneCriterion | error |  | Surrogate | off |
| MaxNumSplits | *n*-1 |  | MinLeafSize | 1 |
| SplitCriterion | gdi |  | OptimizeHyperparameters | none |

where *n* is the number of instances.

③ Random Forest: Number of Decision Trees: 100

④ Naïve Bayes:

| Parameter | Default value |  | Parameter | Default value |
| --- | --- | --- | --- | --- |
| DistributionNames | normal |  | Kernel smoother type | normal |
| Kernel smoothing  density support | unbounded |  | Prior | empirical |
| OptimizeHyperparameters | none |  |  |  |
