## Supplementary Materials for "Information-dense transcription factor binding site clusters identify target genes with similar tissue-wide expression profiles and serve as a buffer against mutations"

**1. Cofactor binding sites flanking YY1 sites in the promoters of the targets and non-targets of YY1**

**Targets:**

① MRPL9:

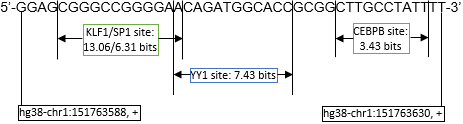

② BAZ1B:

Sequence fragment 1 in the promoter:

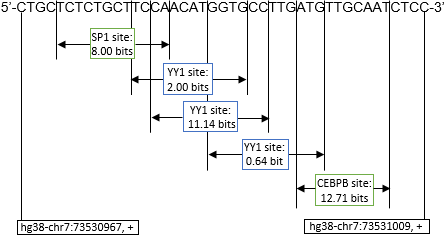

Sequence fragment 2 in the promoter:

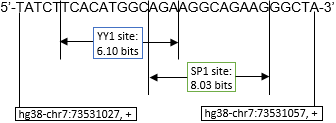

**Non-targets:**

① ADNP:

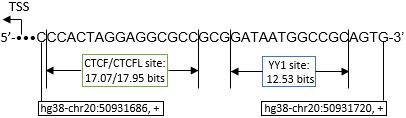

② RNF25:

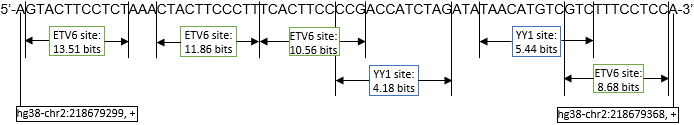

**2. Cofactor binding sites flanking EGR1 sites in the promoters of the targets and non-targets of EGR1**

**Targets:**

① CANX:

Sequence fragment 1 in the promoter:

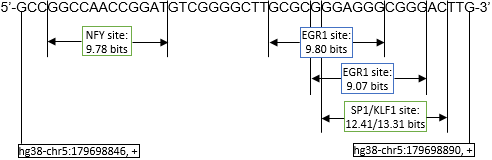

Sequence fragment 2 in the promoter:

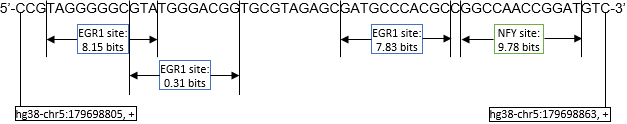

Sequence fragment 3 in the promoter:

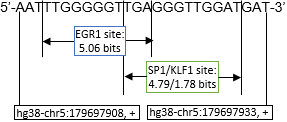

② NPM1:

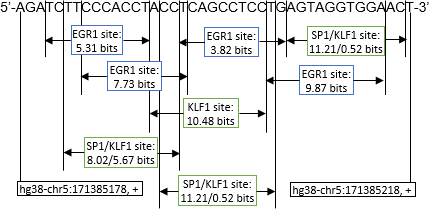

**Non-targets:**

① GUCY2F:

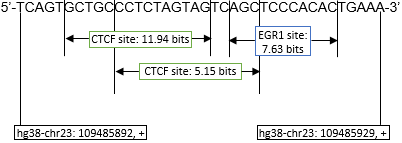

② AWAT1:

Sequence fragment 1 in the promoter:

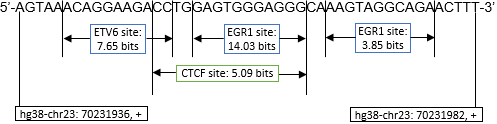

Sequence fragment 2 in the promoter:

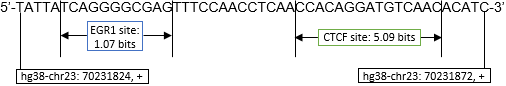
